# Development of a highly active engineered PETase enzyme for polyester degradation

**DOI:** 10.1101/2024.07.04.602061

**Authors:** Shapla Bhattacharya, Rossella Castagna, Hajar Estiri, Toms Upmanis, Andrea Ricci, Alfonso Gautieri, Emilio Parisini

## Abstract

Polyethylene terephthalate (PET) accounts for ≈6% of global plastic production, contributing considerably to the global solid waste stream and environmental plastic pollution. Since the discovery of PET-depolymerizing enzymes, enzymatic PET recycling has been regarded as a promising method for plastic disposal, particularly in the context of a circular economy strategy. However, as the PET degrading enzymes developed so far suffer from relatively limited thermostability, low catalytic efficiency, as well as degradation intermediate-induced inhibition, their large scale industrial applications are still largely hampered. To overcome these limitations, we used *in silico* protein design methods to develop an engineered Leaf-branch Compost Cutinase (LCC), named DRK3, that features enhanced thermal stability and PETase activity relative to the current gold standard LCC enzyme (LCC-ICCG). DRK3 features a 4.1°C increase in melting temperature relative to the LCC-ICCG enzyme. Under optimal reaction conditions (68°C), the DRK3 enzyme hydrolyzes amorphous PET material into TPA with a 2-fold higher efficiency compared to LCC-ICCG. Owing to its enhanced properties, DRK3 may be a promising candidate for future applications in industrial PET recycling processes.

**Graphical abstract:** 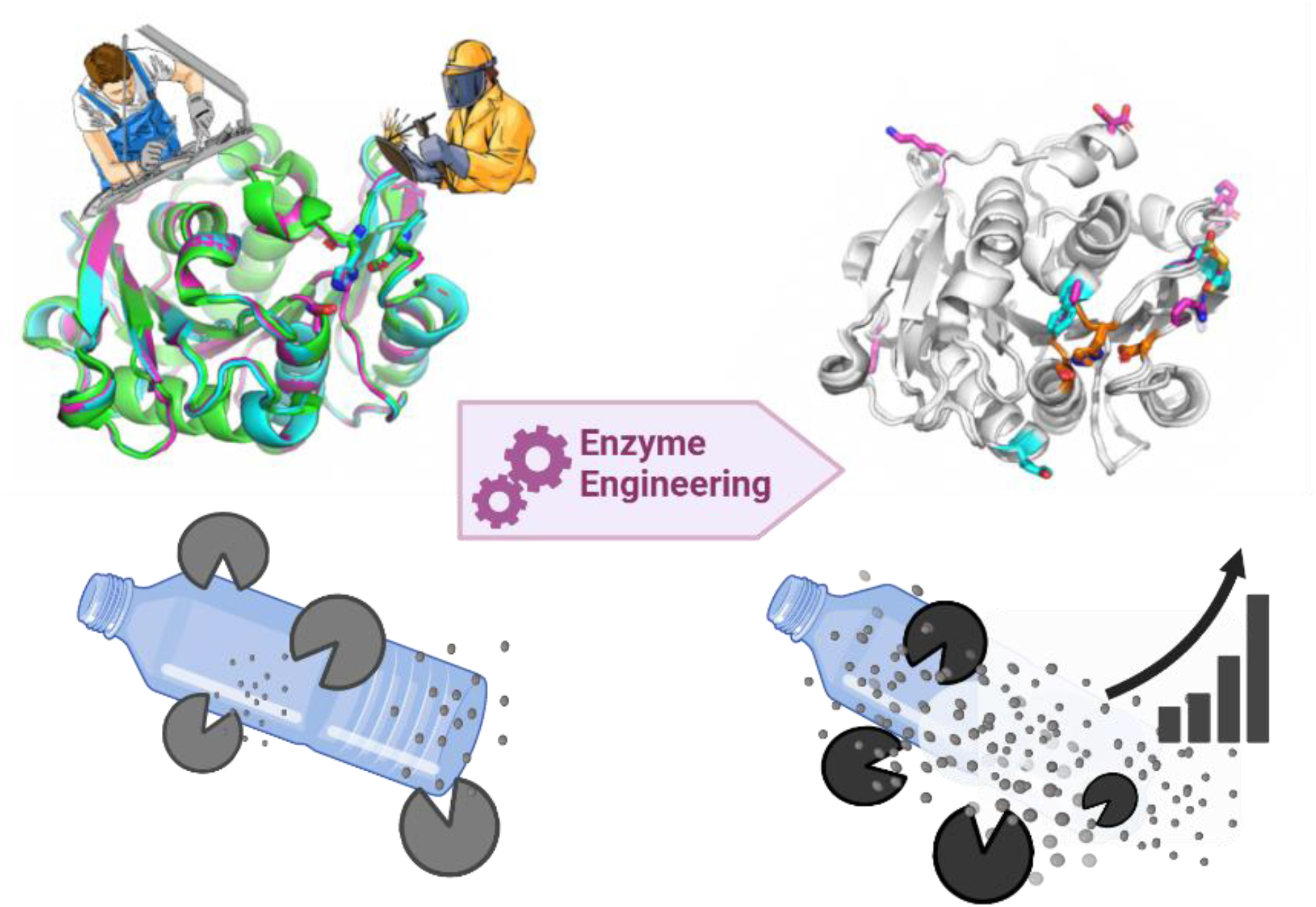

## Introduction

Worldwide plastic production reached 390 million metric tons in 2021, of which 90.2% comes from fossil-based production and only the remaining 9.8% comes from post-consumer plastic recycling or bio-based plastics.^1^ This massive production and usage of plastics, along with their very long persistence in the environment, has led to an extreme global pollution threat, especially in marine environments.^2–5^

Polyethylene terephthalate (PET), the most abundant synthetic polyester in the environment, has an average lifetime of 25–50 years and accounts for 6.2% of the total plastic production.^1^ It is widely used in the production of textile fibers and resins for single-use beverage bottles and packaging. Polyethylene terephthalate is a thermoplastic polymer that is composed of terephthalic acid (TPA) and ethylene glycol (EG) subunits. Nowadays, PET is, to a large extent, mechanically recycled, a process that results in a significant loss of the material’s properties and value.^6^ Harsh chemical treatments involving the use of sulfuric acid at 150 °C or carried out under alkaline conditions in the presence of hazardous chemical catalysts (e.g., methyltrioctylammonium bromide) are also used, leading to the depolymerization of PET into its monomeric building blocks via ester bonds cleavage. However, the use of such harsh conditions makes it problematic to expand this treatment on a large scale.

In 1977, Tokiwa and Suzuki proposed to use lipase enzymes to degrade polymeric materials.^7^ Indeed, enzymes work in mild conditions and can replace hazardous chemicals, a concept known as green chemistry. The first report of an efficient PET hydrolase (from *Thermobifida fusca*) was published in 2005.^8^ Since then, numerous thermally stable PET hydrolases and their related enzymes from the cutinase group (EC. 3.1.1.74) have been identified in different organisms^9–13^. The search for thermostable PET hydrolases is driven by the fact that PET can be more effectively hydrolyzed at temperatures close to its glass transition temperature (≈70-80°C in air, ≈60-70°C in water). Near the glass transition temperature, the polymer chains become more flexible, enabling PET hydrolases to function optimally. In 2016, Yoshida et al.^14^ reported on a mesophilic bacterium (*Ideonella sakaiensis*) that can thrive on an amorphous PET film as its primary carbon source already at 30°C, making the enzyme responsible for PET hydrolysis (*Is*PETase) the best option for PET waste decomposition. This work spurred a great deal of interest and several efforts in providing enhanced *Is*PETase mutants. However, since *Is*PETase is heat-labile, the enzyme quickly loses its activity at temperatures above 40°C. Therefore, more suitable scaffolds have also been explored, such as for instance the Leaf-branch Compost Cutinase (LCC), a naturally occurring PETase that has been reported to outperform all other known PET-degrading enzymes and to present a melting temperature (T_m_) of 84.7°C. This enzyme has been noticeably engineered in 2020 by Tournier et al.,^15^ leading to the so-called ICCG variant (also known as LCC-ICCG), which, in two different accounts, is reported to feature a T_m_ of 91.7 or 94.0°C (Table 1).

**Table 1.** Relevant PETase enzymes reported in the literature.

| Parent Enzyme | Name | Year | Melting Temperature ($^\circ\text{C}$) | References |
| --- | --- | --- | --- | --- |
| IsPETase | (wild type) | 2016 | 46.0 | Yoshida et al.<sup>14</sup> |
| IsPETase | IsPETase<sup>TM</sup> | 2019 | 57.6 | Son et al.<sup>16</sup> |
| LCC | (wild type) | 2020 | 84.7 | Tournier et al.<sup>15</sup> |
| LCC | LCC-ICCG | 2020 | 94.0 | Tournier et al.<sup>15</sup> |
| LCC | LCC-ICCG | 2023 | 91.7 | Arnal et al.<sup>17</sup> |
| IsPETase | - | 2021 | 61.2 | Meng et al.<sup>18</sup> |
| IsPETase | IsPETase<sup>TM</sup><sup>K95N/F201I</sup> | 2021 | 61.6 | Brott et al.<sup>19</sup> |
| IsPETase | DuraPETase | 2021 | 77.0 | Cui et al.<sup>20</sup> |
| IsPETase | HotPETase | 2022 | 82.5 | Bell et al.<sup>21</sup> |
| PES-H1 | L92F/Q94F | 2022 | 86.7 | Pfaff et al.<sup>22</sup> |
| IsPETase | FastPETase | 2023 | 67.1 | Lu et al.<sup>23</sup> |
| IsPETase | DepoPETase | 2023 | 69.4 | Shi et al.<sup>24</sup> |

To the best of our knowledge, this variant is currently considered to be the gold standard PETase enzyme. Despite its remarkable stability, the sequence space for even a small enzyme like LCC is astounding (20^293^ or 10^381^ theoretical possible sequences), leaving room to search for even more stable and active enzymes. Here, we set out to design improved PETase enzymes starting from the LCC-ICCG variant and using design strategies similar to those that we used previously with other classes of enzymes.^25–28^ Our working hypothesis is that improving the thermal stability would be beneficial for the PETase enzymatic activity, as observed in notable examples (IsPETase < LCC < LCC-ICCG). For this reason, we set out to improve LCC-ICCG thermal stability, expecting further beneficial effects such as for instance higher expression yield and enhanced enzymatic activity.

The field of computational design of thermostable proteins is relatively mature, with several established available methods that can help to generate a library of mutants and rank them. The easiest and more common approach involves the introduction of one or more disulphide bonds, which can rigidify the backbone of the protein and make it more stable. Other stabilization methods generally aim at extending the intramolecular hydrogen bonding network, creating inter-molecular salt-bridges, or providing a better packing of the hydrophobic core (e.g., PROSS).^29^ The Rosetta Supercharge method is meant to improve the stability of the protein by making the surface more hydrophilic, a feature that is usually associated with more stable proteins.^30^

While these methods are fast and have been successfully used, most of them use an implicit solvent model (i.e., the solvent is modelled with a continuum rather than with explicit atoms) and assume a rigid backbone (which may limit the long range structural effects of a mutation). To overcome these limitations, we included a further computational filter based on Molecular Dynamics simulations in explicit water. By using an orthogonal method, we aimed at screening and ranking on a common platform the most promising mutants generated by the methods described above.

Our efforts led to the development of three variants that feature T_m_ values higher than any other PETase known to date and, for one of them (DRK3), further enhanced catalytic activity.

## Results and Discussion

### *In silico* design and selection of stabilizing mutants

In the first stage of our *in silico* enzyme engineering campaign, starting from the LCC-ICCG crystal structure (PDB code 6THS) we used three different protein design tools to generate a library of potentially stabilized enzyme variants. The use of the Rosetta Supercharge method,^30^ which aimed at increasing the charge on the surface of the enzyme, led to the design of 1000 enzyme variants, from which those 10 that featured the best Rosetta score (which we named C01-C10) were selected for further analysis. PROSS design^29^ produced 9 variants (P01-P09), and disulfide bond design^31^ identified 125 enzyme variants, from which we selected the best 10, named X01-X10.

The 29 candidate enzyme variants originating from the different screening methods were then subject to 1 μs molecular dynamics (MD) simulations in explicit water and ranked based on their stability as assessed by RMSF analysis.^25^ Our MD-based ranking led to the selection of six candidate mutants (C08, C09, P06, P08, X05, X09) for experimental production and characterization (Figure 1, Figure S1 and Table 2).

**Figure 1.**
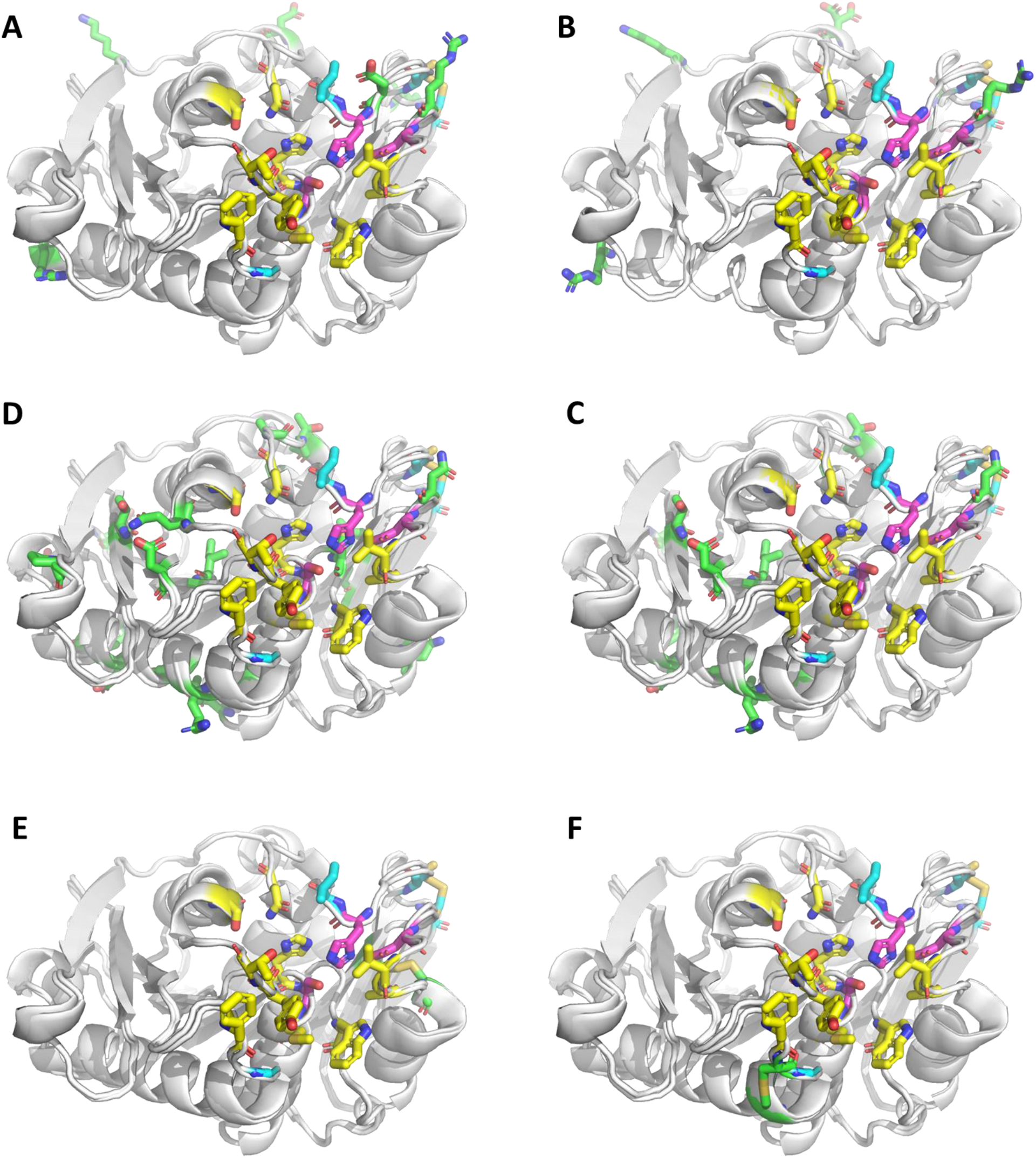
Structural comparison of the enzyme variants selected for experimental characterization. Pink sticks represent the catalytic triad, yellow stick represent amino acid relevant for substrate binding which are prevented from mutations, cyan stick represents the four mutations introduced in LCC-ICCG and green sticks represent additional mutations introduced in each new variant (A: C08, B: C09, C: P06, D: P08, E: X05, F:X09).

**Table 2.** List of mutations relative to the sequence of the wild type LCC enzyme. In bold the mutations already present in the LCC-ICCG scaffold.

| Enzyme | Number of mutations | Mutations |
| --- | --- | --- |
| LCC-ICCG<sup>15</sup> | 4 | Y127G, D238C, F243I, S283C |
| C08 | 11 | S48D, S57K, **Y127G**, S145R, A209R, **D238C**, S241D, **F243I**, N276D, N278K, **S283C** |
| C09 | 13 | S36D, Q40R, S48D, S57K, **Y127G**, S145R, A209R, **D238C**, **F243I**, N276D, N278K, **S283C**, Q293K |
| P06 | 16 | V75T, L84E, M91I, N120D, **Y127G**, S133R, S136L, N140D, A156P, I174A, A209N, **D238C**, **F243I**, A250T, S253A, **S283C** |
| P08 | 26 | S64P, V75T, L84E, M91I, A99Q, N120D, **Y127G**, S133R, S136L, N140D, R143V, A152S, A156P, I174A, N178D, T188C, A209N, Q224N, V233M, V235I, **D238C**, **F243I**, N248S, A250T, S253A, **S283C** |
| X05 | 6 | **Y127G**, G206C, V215C, **D238C**, **F243I**, **S283C** |
| X09 | 6 | D126C, **Y127G**, S130C, **D238C**, **F243I**, **S283C** |

### Thermal stability

The six candidate enzymes were expressed in E.coli and purified. The final yields of all the enzymes were around 16 mg/liter of bacterial culture. Wild type LCC, a naturally occurring PETase, has a T_m_ of 84.7 °C. This enzyme has been previously engineered by Tournier et al.,^15^ leading to the LCC-ICCG variant with a reported T_m_ of 94.0°C (92.9 °C in our own assessment, a difference that could be attributed to that fact that, for T_m_ measurements, Tournier et al. used differential scanning fluorimetry (DSF), whereas we used circular dichroism (CD)). After our enzyme design campaign, we tested the thermal stability of the six engineered designs (Table 3) and we found that the C09 (T_m_ = 97.0 °C) and X05 (T_m_ = 96.2 °C) variants have a higher T_m_ than the reference LCC-ICCG (T_m_ = 92.9 °C). The X09 (T_m_ = 92.3 °C) variant showed a limited improvement in the thermal stability, while C08 (T_m_ = 85.6 °C) showed a slight decrease in the thermal stability (Figure S2-3). For the P06 and P08 variants, we did not set out to determine their melting temperature because of their negligible enzymatic activity (see below).

**Table 3.** Assessed melting temperature for different PETases and improvement with respect to the LCC wild type enzyme. The estimation of T_m_ is sensitive to the experimental setup and conditions, which could explain the difference between the reported T_m_ and the T_m_ determined by us. For proper comparison, we assessed the T_m_ of LCC-ICCG using the same setup that we used for the measurement of the T_m_ of our engineered enzymes. The melting temperature was not assessed for P06 and P08 (n.d., not detected).

| Enzyme | $T_m$ [°C] | Reference |
| --- | --- | --- |
| WT-LCC (reported) | 84.7 | Tournier et al. <sup>15</sup> |
| LCC-ICCG (reported) | 94.0 | Tournier et al. <sup>15</sup> |
| LCC-ICCG (reported) | 91.7 | Arnal et al. <sup>33</sup> |
| LCC-ICCG (assessed) | 92.9±3.6 |  |
| C09 | 97.0±1.0 |  |
| X05 | 96.2±3.7 |  |
| X09 | 92.3±0.6 |  |
| C08 | 85.6±6.1 |  |
| P06 | n.d. |  |
| P08 | n.d. |  |

### Enzymatic activity

Terephthalic acid (TPA) is one of the major degradation products of the activity of the enzymes on PET. This compound can be detected by liquid chromatography (HPLC), which allows a precise quantification of product formation. To assess enzymatic activity, samples were harvested at multiple time points and analyzed by HPLC for the quantification of TPA production (Figure S8). Minor degradation products (MHET and BHET) were also assessed (Figure S9-S10). We used an enzyme concentration of 40 nM and a PET film of 8.4 mg in a 2 ml microplate tube to evaluate the PET degradation activity of the different mutants. The enzyme concentration corresponds to ≈0.3 mg_ENZYME_/g_PET_, a ratio lower than the one reported in the original LCC-ICCG paper (3 mg_ENZYME_/g_PET_), but in agreement with the range reported in the literature (0.2 to 3 mg_ENZYME_/g_PET_).^17,21,32,33^ The experiments were done at 68°C, the temperature at which PET degradation is typically carried out in bioreactors at industrial scale to work below the glass transition temperature of PET (Tg ≈ 70°C). If we exclude the first few hours, where ICCG performs better, at 68°C the mean TPA concentration at the different time points is significantly higher for the C09 mutant than for ICCG up until day 6 (144h, Figure 2A and S5-S8). Moreover, the specific activity of C09 is also significantly higher (≈2-fold) than that of the gold standard LCC-ICCG over the same length of time (Figure 2B). All other enzymes show similar or weaker performances compared to LCC-ICCG (X09, C08, X05). At 68°C, complete sample degradation could be observed with C09 already at day 4 (data not shown), a feature not seen with any of the other enzymes, including LCC-ICCG. The % of weight loss after 6 days of reactions for all the enzymes is shown in Figure 2C and in S11. To assess the stability and the activity of the enzymes in harsh conditions, we also carried out the same experiments at 80 °C for 6 days. Also at this temperature, the mean TPA concentration for the C09 mutant is higher than LCC-ICCG at all the time points (Fig. 2D, S5-S8), albeit being greatly reduced in absolute terms compared to the reaction at 68°C. For the P06 and P08 variants we observed negligible activity (data not shown).

**Figure 2.**
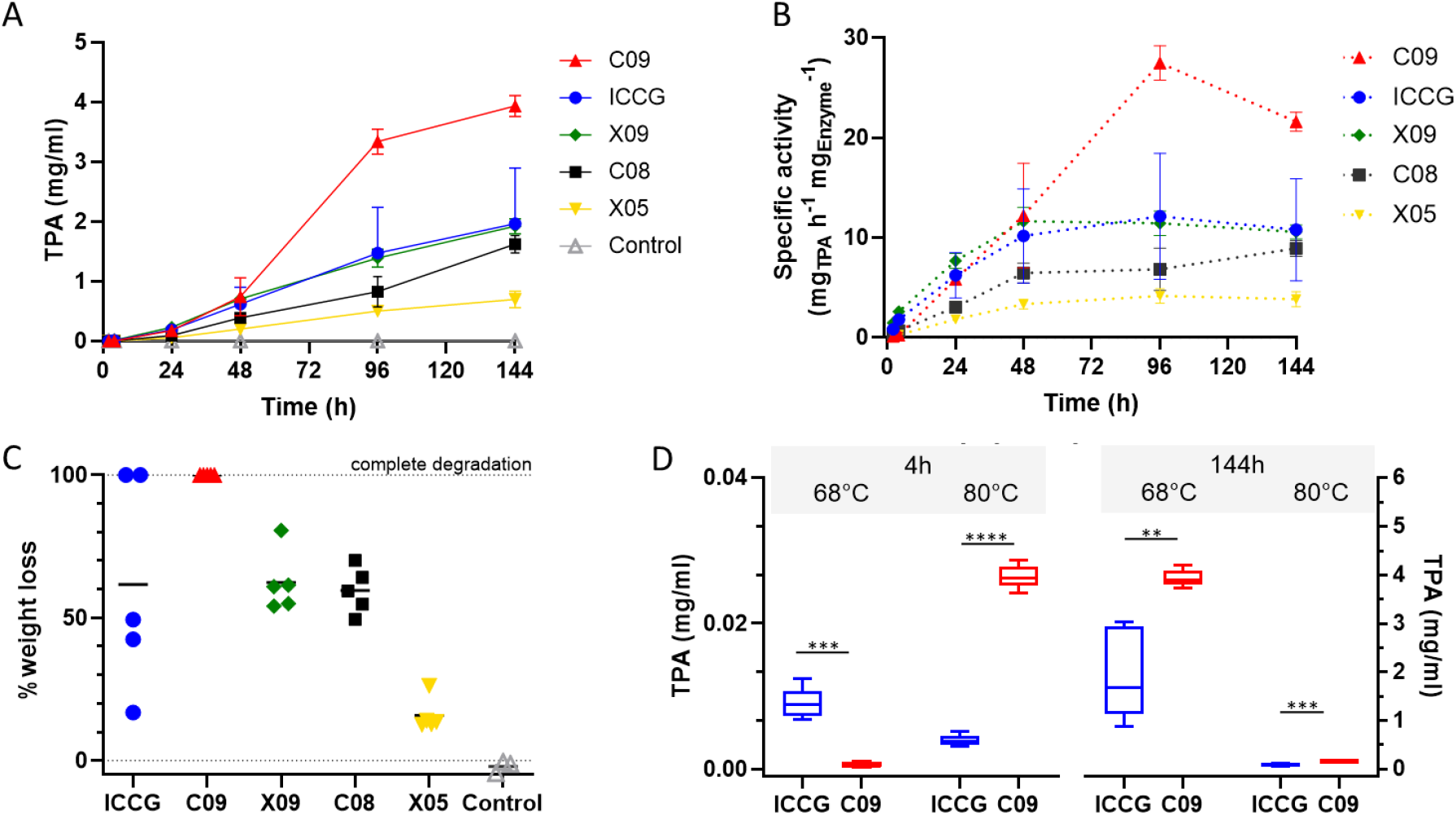
Comparison of PET-depolymerization over time for all enzymes. A) TPA production over time at 68°C (LCC-ICCG – C09 – X05 – X09 – C08 40 nM, pH 8.0). Means ± s.d. (n=5) are shown. B) Enzyme specific activity at the different time points (LCC-ICCG – C09 – X05 – X09 – C08 40 nM, 68°C, pH 8.0). Means ± s.d. (n=5) are shown. C) Percentage of PET film weight loss after treatment (6 days at 68 °C, n=5). The mean % of weight loss (black bar) for LCC-ICCG at day 6 is 61%, while it is 100% for C09. D) Comparison of TPA production at 4h and at 144h, both at 68 and at 80°C. ***P<0.001; ****P<0.0001 (one-tailed unpaired Welch’s t-test). All statistical analyses were performed with GraphPad Prism 10.

### Enzyme structure

We determined the crystal structure of C09 to assess whether the mutations lead to changes in the structure. The structure was determined at 1.28 Å resolution. The structural comparison of the C09 variant with the parent enzymes shows no significant changes in the folding, as observed by the minimal change in RMSD with respect to the WT (0.257 Å) and to the LCC-ICCG variant (0.152 Å). The position and orientation of the catalytic triad (D210, H242, and S265) overlaps perfectly with the catalytic triad in the parent enzymes (Figure 3A). Although we observe that increased stability leads to higher enzymatic activity, a mechanistic explanation of the phenomenon is elusive. We hypothesize that the increase in stability provided by the additional surface charges in the C09 variant may help to keep the surface-exposed catalytic site in place even at high temperatures, such as those used to test the activity of the enzyme (68°C). Indeed, all the mutations introduced in the design process are found mostly on the surface of the enzyme and away from the catalytic triad (Figure 3B), and are thus unlikely to affect the activity directly. Rather, the mutations may reduce the proportion of the enzyme that may denature over time, and/or prevent the unfolding of the exposed catalytic triad. The denaturation (and the consequent loss of activity) is expected to be negligible only for T << T_m_. Therefore, by increasing the T_m_ to 98°C the denaturation effects may be significantly reduced at the working temperature (68°C), explaining the gains in activity of the C09 design. The stabilization of the catalytic triad is confirmed by MD simulations, which show that at increasing temperatures C09 features a lower RMSF, in particular near the key catalytic residue H242 (Figure 3C).

**Figure 3.**
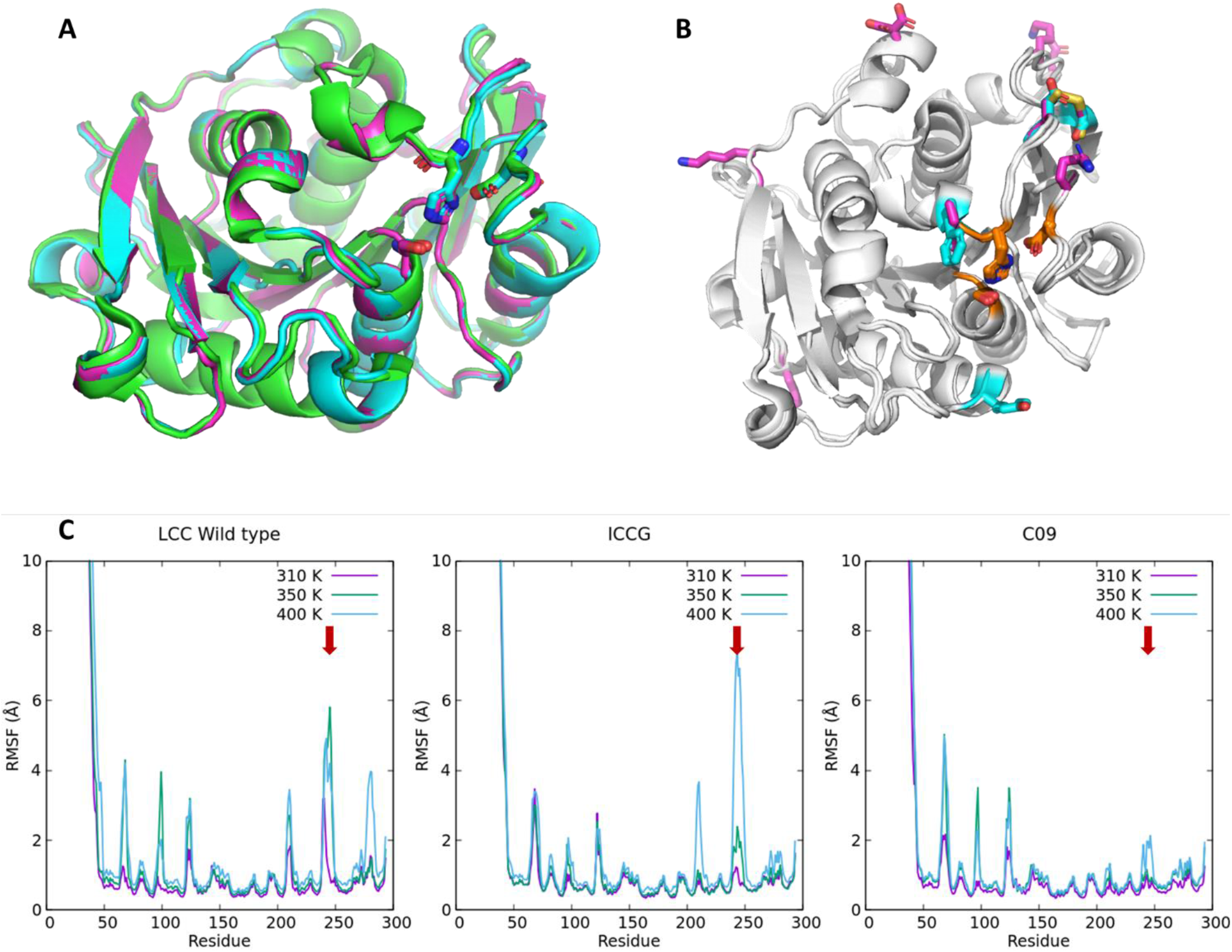
Structural comparison of relevant PETase enzymes: wild type LCC (green), ICCG variant (cyan) and C09 variant (pink). The overall structural alignment shows no significant differences between the enzymes, including the position and the orientation of the catalytic triad (A). Panel B shows the position of mutations (sticks) in ICCG (cyan) and C09 (pink) with respect to the catalytic site (orange sticks). Panel C shows a comparison between the RMSF of LCC, LCC-ICCG, and C09 at different temperatures (310K, 350K and 400k). The RMSF profiles show that the mutations introduced in the C09 variant reduce the structural fluctuations, in particular near the catalytic residue H242 (red arrows).

## Conclusions

Despite *Is*PETase being initially considered a promising enzyme for PET degradation, multiple studies have shown that thermophilic PET hydrolases (e.g., LCC), when operating at temperatures above 65°C, outperform mesophilic enzymes in PET biorecycling.^34^ Since its publication in 2020, the LCC-ICCG variant has been considered the state-of-the-art PETase enzyme. Hence, starting from this variant, we set out to engineer a novel enzyme with enhanced thermal stability and enzymatic activity. The resulting C09 design featured a significant improvement in terms of thermal stability (T_m_=97.0°C) with respect to the gold standard LCC-ICCG (+4.1°C). Under optimal reaction conditions (68°C), the C09 design enzyme hydrolyzed low-crystallinity PET materials into TPA with a 2-fold higher efficiency compared to LCC-ICCG. We nicknamed this design “DRK3”, in light of the introduced mutations (3 Aspartic acids, 3 Arginines, and 3 Lysines). The DRK3 variant may represent a significant advancement towards PET biorecycling. However, further work will be needed to test this enzyme in industrially relevant conditions. The DRK3 variant could also provide the basis for further rounds of enzyme engineering.

Recent studies reported in the literature adopted similar approaches to ours, introducing mutations to the LCC-ICCG enzyme to boost its performance and reporting different degrees of success.^17,32,35,36^ However, as reported in relevant reviews,^37,38^ direct comparison of enzymatic performances is challenging due to the numerous variables involved in the experimental settings (PET crystallinity, PET form, PET concentration, enzyme concentration, temperature, salt type and concentration, pH, etc.). In this light, only a direct comparison with a recognized gold standard in the same experimental conditions can provide reliable information.

Although the field of PETase engineering is very dynamic and is providing highly optimized enzymes, several issues still need to be thoroughly investigated to improve enzymatic PET degradation.^38^ One of the major challenges is related to the economic viability of enzymatic degradation. Another challenge, more directly related to protein engineering, concerns the acidification of the medium due to the release of TPA, which reduces the performance of the PETase enzymes known to date. Hence, possible directions for enzyme engineering research could be directed towards the design of acid-tolerant PETase enzymes.

## Material and methods

### *In silico* design and selection of stabilizing mutants

We started our enzyme engineering process from the LCC-ICCG enzyme reported by Tournier et al. ^15^. This engineered enzyme is a mutant of the wild-type leaf-branch compost cutinase (LCC) carrying 4 point mutations: F243I, Y127G, S283C, D238C. The last two mutations are designed to induce the formation of a second disulfide bond, in addition to the native one (C275-C292). In both enzymes, the catalytic site is centered on residue S165. The structure of the LCC-ICCG engineered enzyme is available in the Protein Data Bank (PDB code: 6THS) as the inactivated S165A variant.

In our design strategy we applied protocols used in previous works^25–27^ starting from the LCC-ICCG mutant structure after reverting amino acid 165 from Alanine to the catalytically active Serine. All residues within a distance of 5 Å from S165 were excluded from the design and were therefore left unchanged. The design was done using three separate methods, resulting in three different series of variants.

The first design strategy (C-series) was performed using the Rosetta Supercharge tool.^30^ This tool was employed to reengineer the protein surface with amino acids that present a high net charge, which is reported to prevent aggregation of partially unfolded states. The design produced 1024 enzyme variants, and the best 10 based on the internal score were kept for further ranking.

In the second design strategy (P-series) we used the PROSS web server^29^ to generate 9 different enzyme designs with improved packing, hydrogen bond networks and salt bridge networks.

The third design strategy (X-series) employed the Disulfide-by-Design web server,^31^ a tool for disulphide bond prediction that allows to find pairs of amino acids that can carry cysteine mutations in the correct orientation to form disulphides. A total of 150 pair mutations were generated and, based on the internal scoring system, the best 10 of them were retained for further testing.

Overall, our design campaign produced 29 different enzyme variants that were then evaluated using MD simulations, and then ranked according to the method described in previous works.^25–27^ Briefly, we solvated each molecular model with a 15 Å pad of TIP3P water and we introduced counter ions to neutralize the system charge, resulting in a final simulation box of ≈30.000 atoms. Hydrogen mass repartitioning was applied to allow a time step of 4 fs ^39^. The systems were subjected to 1,000 energy minimization steps and it equilibrated for 1 ns at a pressure of 1 atm and at a temperature of 300K with NAMD software,^40^ using AMBER19SB force field,^41^ non-bonded cut-off of 12 Å, rigid bonds and particle-mesh Ewald long-range electrostatics. During the equilibration simulation, the Cα atoms of the protein were restrained by a 10 kcal mol^−1^ Å^−2^ spring constant. The 1 μs production runs were performed using a NVT ensemble whereby all the parameters (non-bonded cut-off, and PME) were the same as in the equilibration phase. All simulations were run in triplicates. Root Mean Square Deviations (RMSD) were monitored after each MD run to assess structural convergence, while Root Mean Square Fluctuations (RMSF) were calculated to assess the improved stability of the design. As a result, the two best structures from each design series were selected, namely design C08, C09, P06, P08, X05, X09.

### Protein expression and purification

The different codon-optimized nucleotide sequences of the designed PETase mutants were cloned in a pET26b(+) bacterial expression vector (Novagen), together with a C-terminal 6His-tag. All the different mutants were transformed and expressed in *Escherichia coli* BL21 Star (DE3) cells (Invitrogen). A sample of 4 ml of an overnight culture of the selected strain grown in the presence of 50 µg/ml kanamycin were inoculated into 2L of Luria–Bertani broth (LB) at 37°C, supplemented with 50 mg/L kanamycin and grown until OD_600_ = 0.6. The expression was induced with isopropyl 1-thio-β-D-galactopyranoside (IPTG) at a final concentration of 0.1 mM followed by overnight incubation at 18°C and 220 rpm. Cells were collected by centrifugation and resuspended in lysis buffer (20 mM Tris-HCl pH 8.0, 300 mM NaCl) supplemented with 0.2mM protease phenylmethylsulfonyl fluoride (PMSF) and DNAse and then lysed by sonication on ice. After centrifugation (45000 g for 40 min at 4°C), the soluble fraction of the cell lysate was passed through a 0.45-micron filter and then loaded onto a HisTrap HP 5 ml (GE Healthcare) column equilibrated with buffer A (20 mM Tris-HCl pH 8.0, 300mM Nacl and 10mM imidazole). The Ni affinity column was washed using ten column volumes of buffer A. Before elution, five column volumes (CV) of buffer A supplemented with 50mM imidazole were used to remove unspecific proteins from the resin. Then, the bound His-tagged enzyme was eluted with the same buffer supplemented with 500 mM imidazole in a linear gradient. The protein eluted completely with 250mM imidazole. The eluted fraction was loaded onto a PD-10 desalting column pre-equilibrated with buffer C (20 mM Tris-HCl pH 8.0, 300 mM NaCl) for buffer exchange. Buffer exchange was performed simultaneously after the protein was purified. The desired protein was concentrated to ≈2 mg/ml using an Amicon 20 centrifugal filter with a molecular cutoff of 10 KDa, and stored at −80°C. Protein concentration was assessed using a NanoDrop OneC (Thermo Fisher Scientific). Sample purity was evaluated by 12% SDS-PAGE. The final yields of all the enzymes were around 16 mg/liter of bacterial culture.

### Circular dichroism

Circular dichroism (CD) measurements were performed on a Jasco J-1500 spectropolarimeter at 20 °C. CD was performed to detect the secondary structure of different variants and to measure the thermostability of the enzyme. The spectra were determined with the following parameters: continuous scanning mode with scanning speed of 50 nm/min, band width 0.1 nm, data pitch 0.1 nm at 20 °C. 1.0 mm path length quartz cuvette was used for 5 μM samples of different enzymes in 20mM Tris-HCl pH 8.0, 150mM NaCl. Thermal denaturation curves were measured in 1.0 mm path length cuvettes closed with a parafilm on protein solutions. Protein denaturation was induced upon increasing the temperature at a rate of 1 °C/min from 20 °C to 120 °C. The ellipticity at 222 nm was recorded at intervals of 0.2 °C using a 1-nm bandwidth and a response of 10 s. Using the Boltzmann function, the midpoints of the thermal-denaturation curves (T_m_) were determined by fitting the data to a sigmoidal transition curve^42^. Final data are the average of three measurements. The secondary structures of all the different variants were measured at 200-250 nm wavelengths and 20 °C. Ellipticity was plotted versus wavelength with GraphPad Prism 10 software. Each recorded spectrum was the average of three scans (See Supplementary Information Figure S2).

### Enzymatic reaction

Bis(hydroxyethyl)terephthalate (BHET), mono(hydroxyethyl)terephthalate (MHET), terephthalic acid (TPA), and ethylene glycol (EG) are the major depolymerization products of the activity of the enzymes on PET. These compounds can be separated efficiently on a C18 reversed-phase HPLC column, thus allowing a precise quantification of product formation. All enzyme reactions were performed at a concentration of 40 nM at both 68 and 80 °C in quintuplicates in 2.0 mL microcentrifuge tubes in the same reaction buffer (20 mM Tris-HCl pH 8.0, 300mM NaCl). Amorphous, 250 μm-thick PET films (product number ES301445/11) were purchased from Goodfellow USA and cut into 6 mm diameter circular pellets using a hole puncher. Individual pellets, whose weight was around 8.4 mg, were then placed in 2 ml microcentrifuge tubes.

### Reversed-phase liquid chromatography

The different compounds produced by PET hydrolytic enzymes (TPA, as well as BHET and MHET) were quantified by reversed-phase HPLC (See Supporting Information Table S2-S4). HPLC analyses were performed on a Shimadzu LC-2030C 3D Plus system equipped with a PDA detector, a column oven and an autosampler (Shimadzu, Kyoto, Japan). Separations were carried out using a Kinetex C18 column (2.7 µm; 4.6 × 150 mm, Phenomenex, Torrance, CA, USA). Column temperature was maintained at 40°C with a flow rate of 1 ml/min. For the mobile phase, 0.1% H_3_PO_4_ (solvent A) and MeCN (solvent B) were used. The linear gradient mode used for the elution was as follows: 0 min, 10% B; 0-15 min, 10-40% B; 15-18min, 40-65% B; 18-20 min, 65% B; 20-23 min, 65-10% B; 23-25 min 10% B. At every time interval, 100 μL of the sample were collected from the reaction vial placed at the set temperature. The 0h–72h samples were diluted 21 times (50µL of the provided sample was transferred to autosampler vial and diluted in 1 mL of 10% MeCN + 0.1% H_3_PO_4_), whereas, 96h and 144h sampes – 101 times (10µL of the provided sample was transferred to autosampler vial and diluted in 1 mL of 10% MeCN + 0.1% H_3_PO_4_). Detection was accomplished via measurement of UV adsorption at 220 nm; 20 μL of each sample was injected into the system and the degradation products were quantified against standard calibration curves ranged 0.1, 1.0, 5.0, 10.0, 50.0 (for TPA and MHET) and 0.09, 0.9, 4.4 and 43.9 (for BHET) μg/mL.

### Protein crystallization

Crystals of the C09 mutant were grown at room temperature using the vapor diffusion method by mixing a 2μL drop of a 10mg/mL protein sample with a 2μL drop of a solution containing 0.1M Sodium citrate tribasic dihydrate pH 5.6, 20% v/v 2-Propanol, 20% w/v Polyethylene glycol 4,000. Crystals, which appeared after one week, were frozen in liquid nitrogen using 25% (v/v) glycerol as cryoprotectant prior to X-ray diffraction data collection.

### Structure determination and refinement

For the structure of the C09 mutant, the initial phases were obtained by molecular replacement using Phaser^43^ and the atomic coordinates of the crystal structure of a Leaf-branch compost bacterial cutinase homolog (PDB entry 4EB0) as a search model. Refinement was performed by alternating rounds of REFMAC5^44^ and manual adjustments in Coot.^45^ Water molecules were added both manually and automatically using the Coot_refine tool from the CCP4 package.^46^ The crystallographic table is shown in the SI file in Table S7.

## Data availability

The final crystallographic coordinates of the crystal structures of the C09 mutant are available in the RCSB PDB (accession code: 8CMV).

## Supporting Information

Additional experimental details including sequence alignment, data on circular dichroism, enzymatic activity and crystallographic data.

## Acknowledgements

The authors wish to thank the personnel of the Diamond Light Source (proposal number mx32544) for help with the data collection. E.P. thanks the European Regional Development Fund (ERDF) project BioDrug (No. 1.1.1.5/19/A/004), the Latvian Council of Science and the Latvian Recovery and Resilience Fund (grant No. lzp-2020/2-0013 and grant No. 74/OSI/ZG) for financial support. S.B. acknowledges the Latvian Recovery and Resilience Fund and the Latvian Institute of Organic Synthesis for student grants (No. ANM_OSI_DG_12 and No. IG-2024-08). R.C. acknowledges the MICS (Made in Italy — Circular and Sustainable) Extended Partnership and received funding from Next-Generation EU (Italian PNRR – M 4 C2, Invest 1.3 – D.D. 1551.11–10-2022, PE00000004) CUP MICS C93C220052800. E.P. and A.G. wish to thank the European Union’s HORIZON-WIDERA-2023-ACCESS-04 programme for financial support under grant agreement 101159534 (WIDEnzymes). This manuscript reflects only the authors’ views and opinions. Neither the European Union nor the granting authority can be considered responsible for them. We acknowledge the CINECA award under the ISCRA initiative (Grant code IsB26_W2EB and IsCa2_REZYME) for the availability of high-performance computing resources and support. Graphs were done with GraphPad Prism 10 software and the graphical abstract was created with BioRender.com.

## Contributions

Conceptualization: AG, EP; methodology: SB, HE, TU, RC, AG, EP; formal analysis: SB, HE, AR, RC; investigation: SB, HE, TU, AR, RC; data curation: SB, RC, AG, EP; writing, review and editing: SB, HE, AR, RC, AG, EP; supervision: RC, AG, EP; funding acquisition: AG, EP. All authors have read and agreed to the published version of the manuscript.

