## Supplementary data for "Development of a highly active engineered PETase enzyme for polyester degradation"

Number of Pages: 15
Number of Figures: 11
Number of Tables: 7

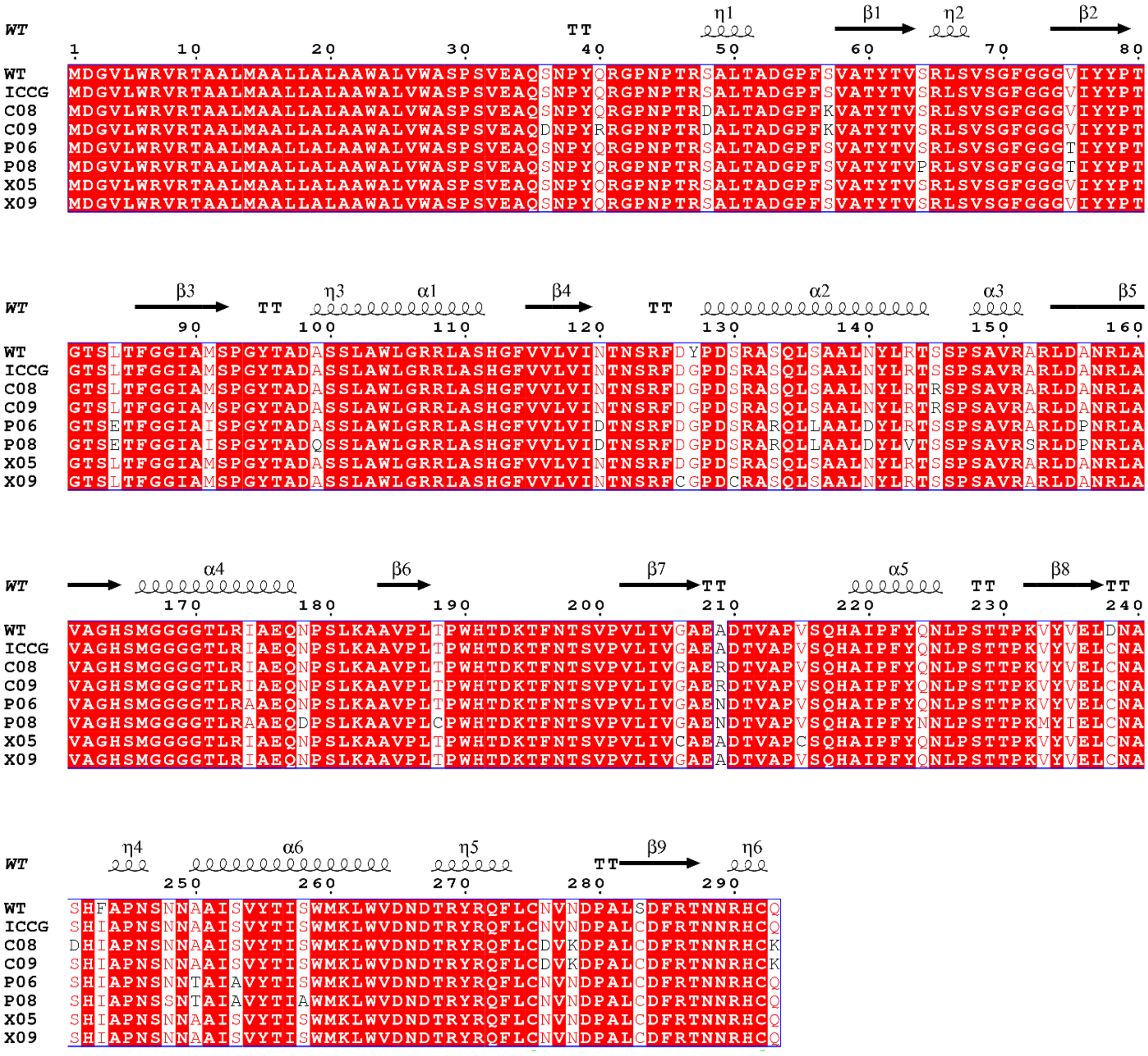

**Figure S1.** Sequence alignment of the wild type LCC enzyme (Uniprot ID G9BY57), the ICCG variant and the engineered enzymes developed in this work.

Circular dichroism curves corresponding to the denaturation temperature (Tm) for the different mutants, measured from 20 °C to 120 °C at 222 nm.

The determination of the midpoints of the thermal-denaturation curves (Tm) was done by fitting the data to a sigmoidal transition curve using the Boltzmann function with the Thermal Denaturation Analysis software of Jasco J-1500.

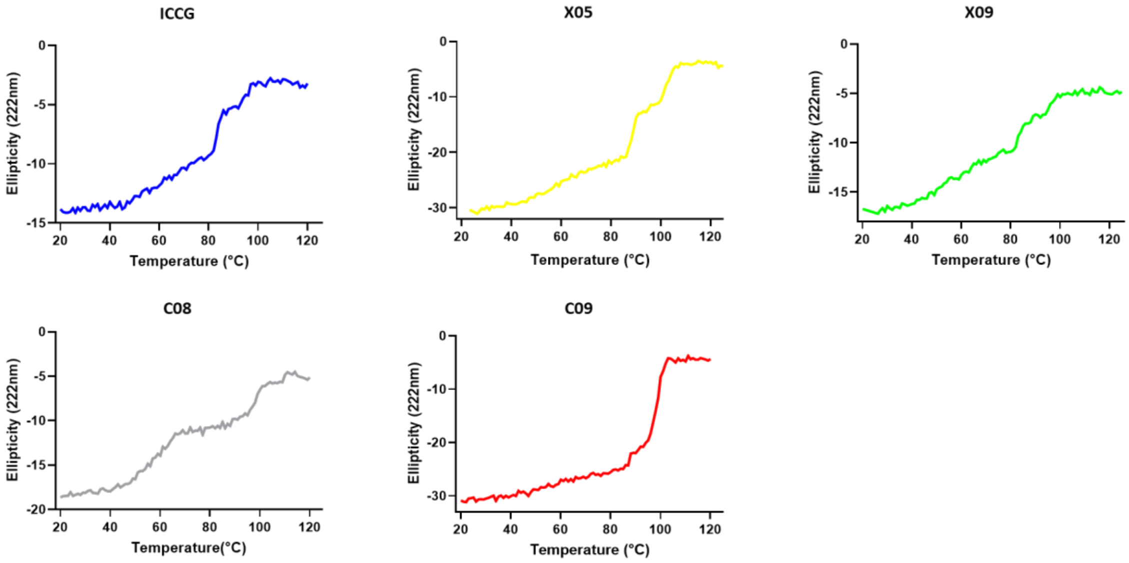


**Figure S2.** Representative melting curves. Tm of the different enzyme variants as measured by circular dichroism at 222nm from 20°C to 120°C.

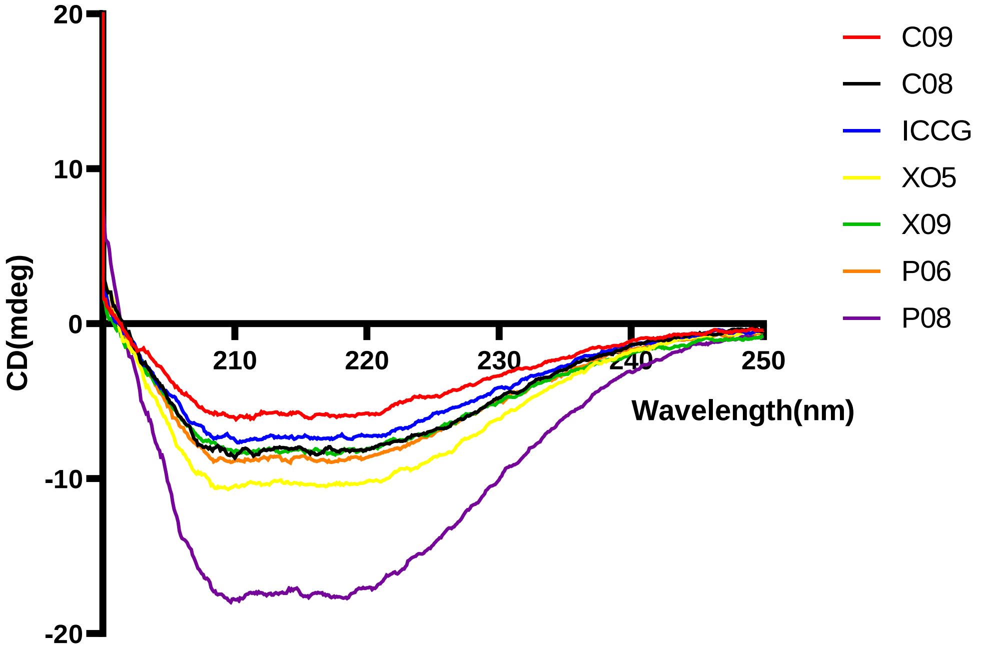


**Figure S3.** Secondary structure measurements by circular dichroism: ellipticity vs. wavelength.

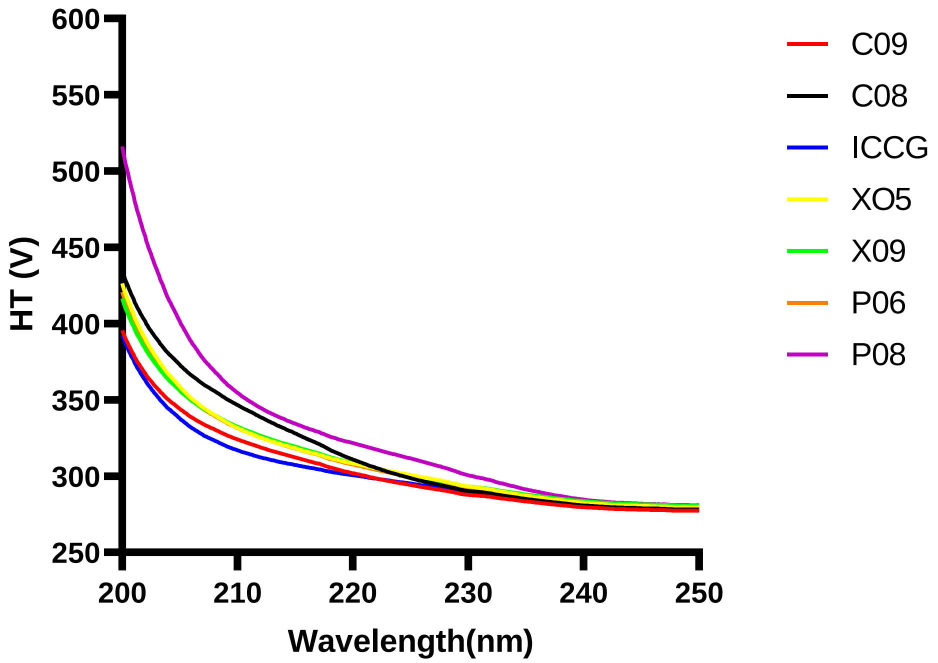


**Figure S4.** Secondary structure measurements by circular dichroism: voltage vs. wavelength.

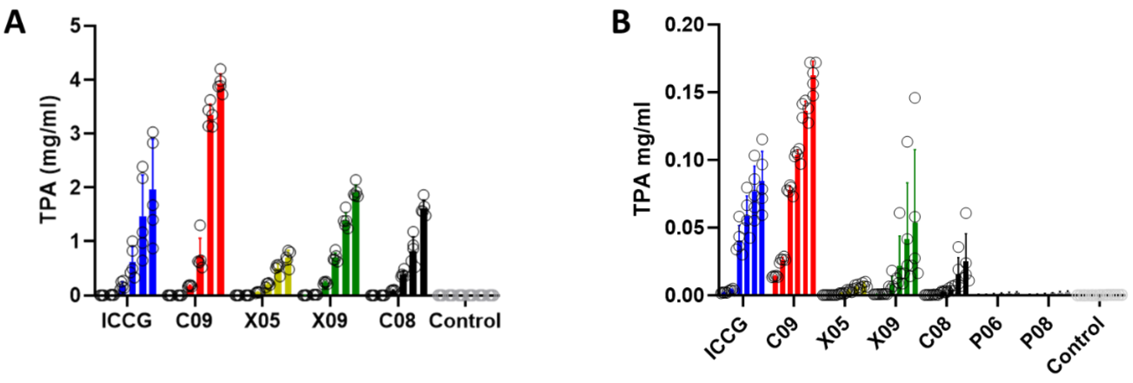


**Figure S5**. Graph of the time course of the TPA production from different enzymes. (A) Experiments performed using 40 nM enzyme at 68°C (n=5). Time points: 2h, 4h, 24h, 48h, 96h and 144h. (B) Experiments performed using 40 nM enzyme at 80°C. Time points: 2h, 4h, 24h, 48h, 96h and 144h (n=5).

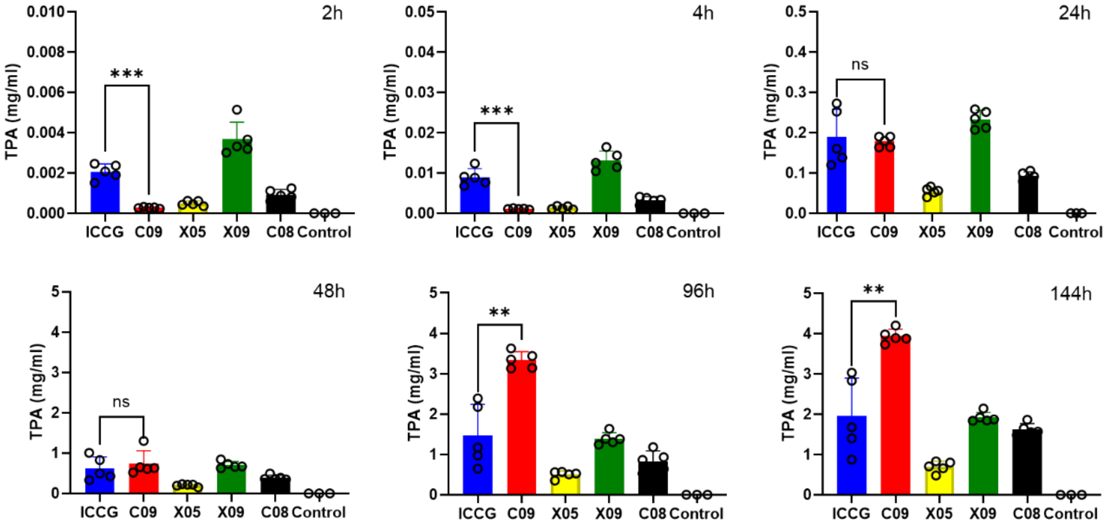


**Figure S6**. TPA production from different enzymes (ICCG – C09 – X05 – X09 – C08 (n=5)) and control (no enzyme (n=3)) at 40 nM, 68°C and at different time points (2h, 4h, 24h, 48h, 96h and 144h). ***P<0.001; ****P<0.0001 (one-tailed unpaired Welch's t-test).

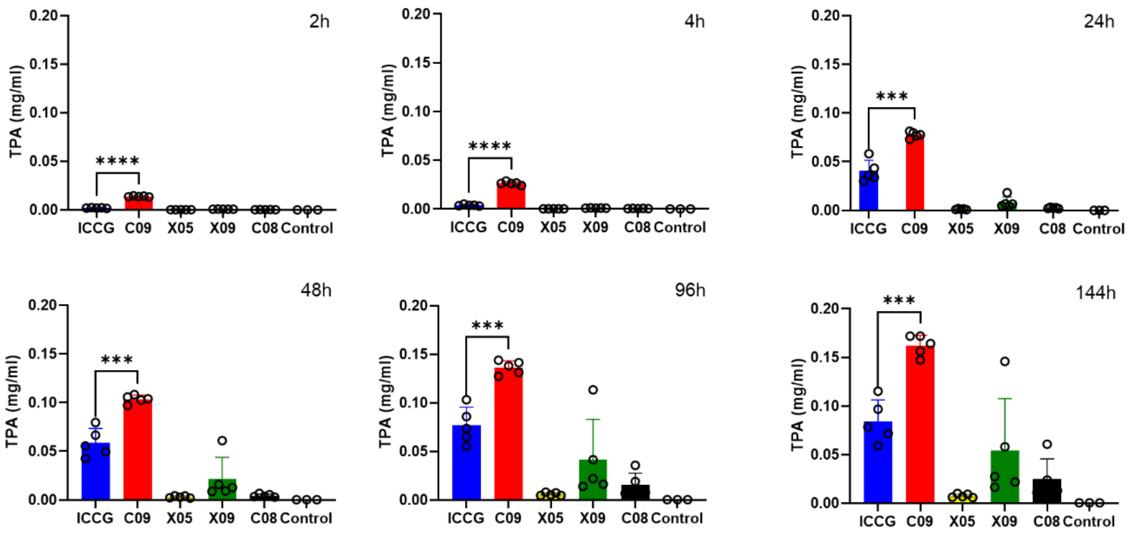


**Figure S7**. TPA production from different enzymes (ICCG – C09 – X05 – X09 – C08 (n=5)) and control (no enzyme (n=3)) at 40 nM, 80°C and at different time points (2h, 4h, 24h, 48h, 96h and 144h). ***P<0.001; ****P<0.0001 (one-tailed unpaired Welch's t-test).

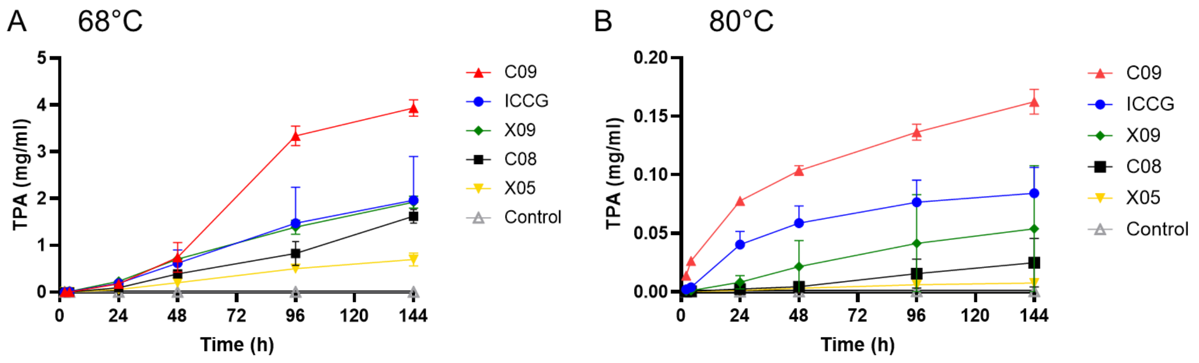


**Figure S8**. **TPA** production over time quantified by HPLC analysis after reaction with 40 nM enzyme (ICCG – C09 – X05 – X09 – C08 (n=5) – no enzyme (control n=3)). A) Experiments performed at 68°C. B) Experiments performed at 80°C.

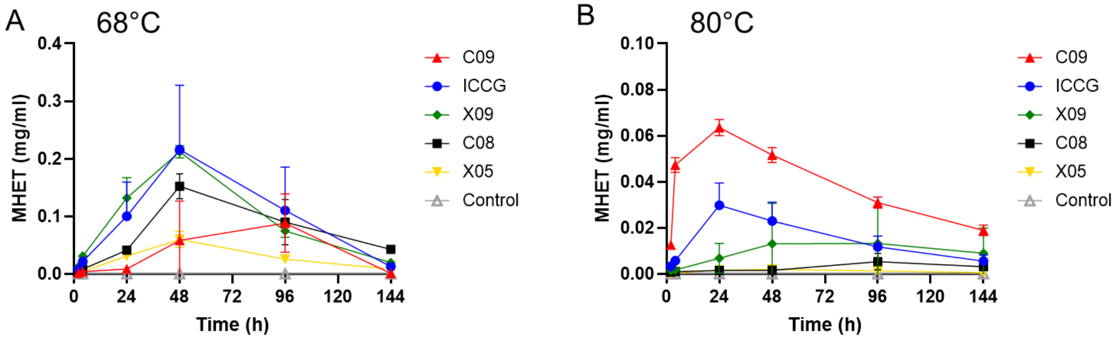


**Figure S9**. **MHET** production over time quantified by HPLC analysis after reaction with 40 nM enzyme (ICCG – C09 – X05 – X09 – C08 (n=5) – no enzyme (control n=3)). A) Experiments performed at 68°C. B) Experiments performed at 80°C.

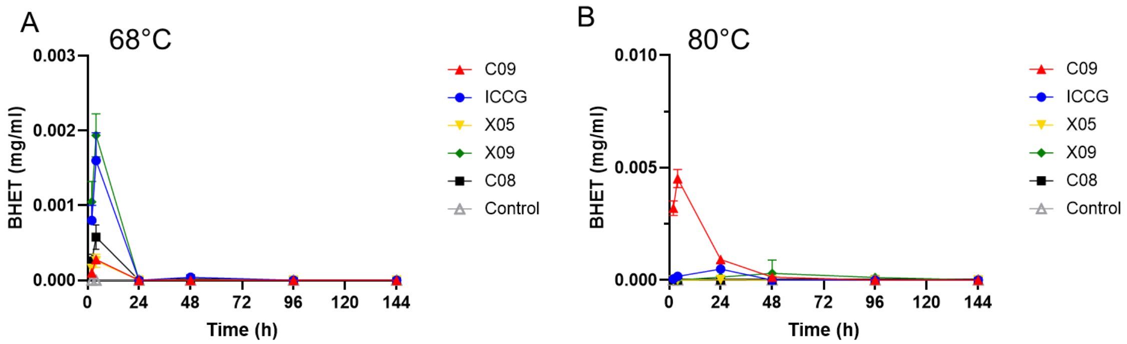


**Figure S10. BHET** production over time quantified by HPLC analysis after reaction with 40 nM enzyme (ICCG – C09 – X05 – X09 – C08 (n=5) - no enzyme (control n=3)). A) Experiments performed at 68°C. B) Experiments performed at 80°C.

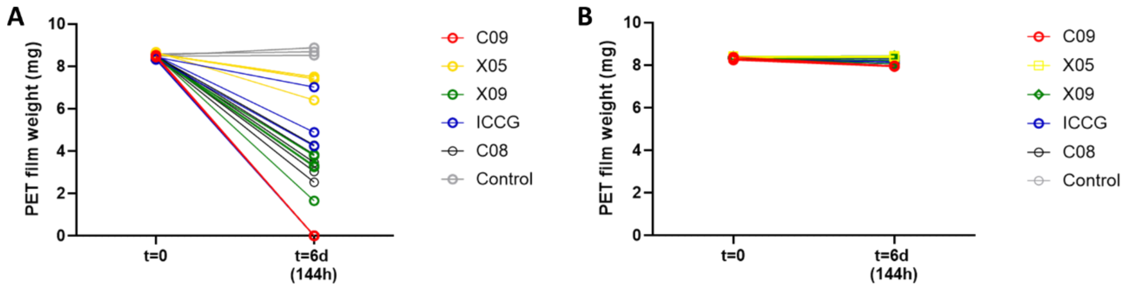


**Figure S11.** PET film weight before (t=0) and after treatment (t=6) with 40 nM enzyme (ICCG – C09 – X05 – X09 – C08 (n=5)). (A) Experiments performed at 68°C. Mean weight loss for ICCG at 6 days is 61%. Mean weight loss for C09 at 6 days is 100%. (B) Experiments performed at 80°C. Mean weight loss for ICCG at 6 days is 1.92%. Mean weight loss for C09 at 6 days is 4.21%.

**Table S1.** HPLC quantification of **TPA** (mg/mL) in reaction supernatants represented in Figure 2A. Reactions were carried out using 40 nM enzyme at 68°C.

| **Enzyme (replicate #)** | **Time, h** | | | | | |
|---|---|---|---|---|---|---|
| | **2** | **4** | **24** | **48** | **96** | **144** |
| **C09(1)** | 0.00023 | 0.0009 | 0.16 | 0.52 | 3.14 | 3.73 |
| **C09(2)** | 0.00028 | 0.0011 | 0.16 | 1.30 | 3.63 | 3.88 |
| **C09(3)** | 0.00028 | 0.0011 | 0.18 | 0.65 | 3.35 | 4.20 |
| **C09(4)** | 0.00032 | 0.0013 | 0.19 | 0.63 | 3.44 | 3.98 |
| **C09(5)** | 0.00026 | 0.0012 | 0.19 | 0.61 | 3.13 | 3.89 |
| **X05(1)** | 0.00043 | 0.0011 | 0.060 | 0.23 | 0.57 | 0.83 |
| **X05(2)** | 0.00062 | 0.0019 | 0.056 | 0.22 | 0.56 | 0.80 |
| **X05(3)** | 0.00046 | 0.0012 | 0.052 | 0.20 | 0.50 | 0.66 |
| **X05(4)** | 0.00061 | 0.0018 | 0.066 | 0.22 | 0.53 | 0.71 |
| **X05(5)** | 0.00035 | 0.0010 | 0.040 | 0.15 | 0.35 | 0.48 |
| **X09(1)** | 0.0033 | 0.0125 | 0.26 | 0.84 | 1.64 | 2.14 |
| **X09(2)** | 0.0032 | 0.0115 | 0.21 | 0.66 | 1.38 | 1.84 |
| **X09(3)** | 0.0051 | 0.0165 | 0.25 | 0.73 | 1.24 | 1.92 |
| **X09(4)** | 0.0030 | 0.0105 | 0.21 | 0.63 | 1.30 | 1.85 |
| **X09(5)** | 0.0037 | 0.0144 | 0.23 | 0.67 | 1.38 | 1.87 |
| **ICCG(1)** | 0.0025 | 0.0124 | 0.25 | 0.83 | 2.18 | 2.83 |
| **ICCG(2)** | 0.0021 | 0.0091 | 0.27 | 1.01 | 2.39 | 3.03 |
| **ICCG(3)** | 0.0024 | 0.0089 | 0.16 | 0.50 | 1.17 | 1.68 |
| **ICCG(4)** | 0.0019 | 0.0077 | 0.12 | 0.33 | 0.65 | 0.87 |
| **ICCG(5)** | 0.0015 | 0.0068 | 0.14 | 0.43 | 0.97 | 1.40 |
| **C08(1)** | 0.0009 | 0.0020 | 0.079 | 0.35 | 1.18 | 1.87 |
| **C08(2)** | 0.0008 | 0.0034 | 0.084 | 0.36 | 0.93 | 1.57 |
| **C08(3)** | 0.0012 | 0.0042 | 0.091 | 0.50 | 0.86 | 1.48 |
| **C08(4)** | 0.0011 | 0.0039 | 0.106 | 0.39 | 0.64 | 1.65 |
| **C08(5)** | 0.0006 | 0.0032 | 0.099 | 0.36 | 0.53 | 1.56 |
| **Control(1)** | < LOD | <LOD | < LOD | <LOD | < LOD | <LOD |
| **Control(2)** | < LOD | <LOD | < LOD | <LOD | < LOD | <LOD |
| **Control(3)** | < LOD | <LOD | < LOD | <LOD | < LOD | <LOD |

< LOQ - below the lowest analyte concentration that can be quantitatively detected with a stated accuracy and precision;
< LOD - no peak found

**Table S2.** HPLC quantification of **TPA** (mg/mL) in reaction supernatants represented in Figure S8B. Reactions were carried out using 40 nM enzyme at 80°C.

| **Enzyme** | **Time, h** | | | | | |
|---|---|---|---|---|---|---|
| **(replicate #)** | **2** | **4** | **24** | **48** | **96** | **144** |
| **C09(1)** | 0.0146 | 0.0287 | 0.0798 | 0.1058 | 0.1313 | 0.1643 |
| **C09(2)** | 0.0136 | 0.0262 | 0.0731 | 0.0971 | 0.1275 | 0.1474 |
| **C09(3)** | 0.0137 | 0.0262 | 0.0766 | 0.1026 | 0.1380 | 0.1565 |
| **C09(4)** | 0.0135 | 0.0267 | 0.0777 | 0.1040 | 0.1414 | 0.1719 |
| **C09(5)** | 0.0143 | 0.0241 | 0.0815 | 0.1083 | 0.1439 | 0.1720 |
| **X05(1)** | 0.00022 | 0.00029 | 0.00198 | 0.00422 | 0.00737 | 0.00918 |
| **X05(2)** | 0.00019 | 0.00024 | 0.00079 | 0.00196 | 0.00467 | 0.00596 |
| **X05(3)** | 0.00019 | 0.00022 | 0.00100 | 0.00227 | 0.00477 | 0.00620 |
| **X05(4)** | 0.00019 | 0.00020 | 0.00116 | 0.00268 | 0.00545 | 0.00673 |
| **X05(5)** | 0.00024 | 0.00024 | 0.00158 | 0.00408 | 0.00804 | 0.01011 |
| **X09(1)** | 0.0008 | 0.0012 | 0.0183 | 0.0610 | 0.1134 | 0.1459 |
| **X09(2)** | 0.0007 | 0.0010 | 0.0067 | 0.0159 | 0.0416 | 0.0580 |
| **X09(3)** | 0.0008 | 0.0010 | 0.0047 | 0.0088 | 0.0141 | 0.0164 |
| **X09(4)** | 0.0007 | 0.0009 | 0.0048 | 0.0092 | 0.0163 | 0.0219 |
| **X09(5)** | 0.0009 | 0.0011 | 0.0066 | 0.0128 | 0.0221 | 0.0275 |
| **ICCG(1)** | 0.0025 | 0.0051 | 0.0583 | 0.0796 | 0.1033 | 0.1153 |
| **ICCG(2)** | 0.0020 | 0.0038 | 0.0435 | 0.0666 | 0.0859 | 0.0968 |
| **ICCG(3)** | 0.0022 | 0.0039 | 0.0301 | 0.0424 | 0.0552 | 0.0593 |
| **ICCG(4)** | 0.0016 | 0.0031 | 0.0339 | 0.0495 | 0.0651 | 0.0717 |
| **ICCG(5)** | 0.0019 | 0.0035 | 0.0363 | 0.0554 | 0.0738 | 0.0784 |
| **C08(1)** | 0.0004 | 0.0007 | 0.0030 | 0.0055 | 0.0191 | 0.0240 |
| **C08(2)** | 0.0004 | 0.0007 | 0.0034 | 0.0068 | 0.0357 | 0.0608 |
| **C08(3)** | 0.0003 | 0.0004 | 0.0022 | 0.0036 | 0.0077 | 0.0131 |
| **C08(4)** | 0.0003 | 0.0005 | 0.0021 | 0.0034 | 0.0070 | 0.0109 |
| **C08(5)** | 0.0003 | 0.0004 | 0.0018 | 0.0030 | 0.0084 | 0.0155 |
| **Control(1)** | < LOD | < LOD | 0.0001 | 0.0002 | 0.0004 | 0.0005 |
| **Control(2)** | < LOD | < LOD | 0.0001 | 0.0002 | 0.0004 | 0.0005 |
| **Control(3)** | < LOD | < LOD | 0.0001 | 0.0002 | 0.0004 | 0.0005 |

< LOQ - below the lowest analyte concentration that can be quantitatively detected with a stated accuracy and precision;
< LOD - no peak found

**Table S3.** HPLC quantification of **MHET** (mg/mL) in reaction supernatants. Reactions were carried out using 40 nM enzyme at 68°C.

| **Enzyme (replicate #)** | **Time, h** | | | | | |
|---|---|---|---|---|---|---|
| | **2** | **4** | **24** | **48** | **96** | **144** |
| **C09(1)** | 0.0011 | 0.0032 | 0.008 | 0.02 | 0.14 | 0.0005 |
| **C09(2)** | 0.0013 | 0.0037 | 0.008 | 0.18 | 0.01 | 0.0004 |
| **C09(3)** | 0.0014 | 0.0035 | 0.010 | 0.03 | 0.07 | 0.0003 |
| **C09(4)** | 0.0016 | 0.0041 | 0.009 | 0.03 | 0.12 | 0.0007 |
| **C09(5)** | 0.0014 | 0.0039 | 0.007 | 0.03 | 0.10 | 0.0024 |
| **X05(1)** | 0.0015 | 0.0034 | 0.035 | 0.08 | 0.033 | 0.011 |
| **X05(2)** | 0.0025 | 0.0055 | 0.030 | 0.06 | 0.029 | 0.010 |
| **X05(3)** | 0.0017 | 0.0034 | 0.032 | 0.06 | 0.026 | 0.008 |
| **X05(4)** | 0.0026 | 0.0056 | 0.035 | 0.06 | 0.026 | 0.008 |
| **X05(5)** | 0.0015 | 0.0034 | 0.022 | 0.04 | 0.014 | 0.005 |
| **X09(1)** | 0.011 | 0.030 | 0.19 | 0.22 | 0.093 | 0.023 |
| **X09(2)** | 0.011 | 0.028 | 0.11 | 0.22 | 0.074 | 0.019 |
| **X09(3)** | 0.017 | 0.037 | 0.14 | 0.22 | 0.063 | 0.018 |
| **X09(4)** | 0.010 | 0.025 | 0.10 | 0.20 | 0.073 | 0.020 |
| **X09(5)** | 0.012 | 0.034 | 0.12 | 0.20 | 0.070 | 0.017 |
| **ICCG(1)** | 0.0076 | 0.028 | 0.17 | 0.30 | 0.18 | 0.0101 |
| **ICCG(2)** | 0.0087 | 0.019 | 0.16 | 0.36 | 0.20 | 0.0082 |
| **ICCG(3)** | 0.0090 | 0.022 | 0.06 | 0.19 | 0.08 | 0.0203 |
| **ICCG(4)** | 0.0054 | 0.020 | 0.05 | 0.09 | 0.03 | 0.0063 |
| **ICCG(5)** | 0.0066 | 0.019 | 0.06 | 0.14 | 0.06 | 0.0184 |
| **C08(1)** | 0.0019 | 0.0050 | 0.044 | 0.14 | 0.15 | 0.0472 |
| **C08(2)** | 0.0037 | 0.0087 | 0.034 | 0.15 | 0.10 | 0.0429 |
| **C08(3)** | 0.0041 | 0.0101 | 0.039 | 0.19 | 0.09 | 0.0388 |
| **C08(4)** | 0.0029 | 0.0097 | 0.046 | 0.14 | 0.06 | 0.0434 |
| **C08(5)** | 0.0031 | 0.0081 | 0.043 | 0.14 | 0.05 | 0.0401 |
| **Control(1)** | < LOD | <LOD | < LOD | <LOD | < LOD | <LOD |
| **Control(2)** | < LOD | <LOD | < LOD | <LOD | < LOD | <LOD |
| **Control(3)** | < LOD | <LOD | < LOD | <LOD | < LOD | <LOD |

< LOQ - below the lowest analyte concentration that can be quantitatively detected with a stated accuracy and precision;
< LOD - no peak found

**Table S4.** HPLC quantification of **MHET** (mg/mL) in reaction supernatants. Reactions were carried out using 40 nM enzyme at 80°C.

| **Enzyme (replicate #)** | **Time, h** | | | | | |
|---|---|---|---|---|---|---|
| | **2** | **4** | **24** | **48** | **96** | **144** |
| **C09(1)** | 0.0132 | 0.0516 | 0.0666 | 0.0542 | 0.0312 | 0.0203 |
| **C09(2)** | 0.0123 | 0.0470 | 0.0579 | 0.0463 | 0.0271 | 0.0155 |
| **C09(3)** | 0.0123 | 0.0472 | 0.0623 | 0.0506 | 0.0307 | 0.0176 |
| **C09(4)** | 0.0121 | 0.0479 | 0.0658 | 0.0538 | 0.0335 | 0.0208 |
| **C09(5)** | 0.0129 | 0.0427 | 0.0653 | 0.0531 | 0.0325 | 0.0201 |
| **X05(1)** | 0.00005 | 0.00055 | 0.00271 | 0.00265 | 0.00139 | 0.00061 |
| **X05(2)** | 0.00002 | 0.00038 | 0.00102 | 0.00130 | 0.00108 | 0.00053 |
| **X05(3)** | 0.00003 | 0.00046 | 0.00114 | 0.00156 | 0.00106 | 0.00047 |
| **X05(4)** | 0.00003 | 0.00047 | 0.00140 | 0.00181 | 0.00111 | 0.00050 |
| **X05(5)** | 0.00007 | 0.00055 | 0.00182 | 0.00250 | 0.00165 | 0.00077 |
| **X09(1)** | 0.0015 | 0.0020 | 0.0183 | 0.0442 | 0.0399 | 0.0275 |
| **X09(2)** | 0.0013 | 0.0017 | 0.0050 | 0.0090 | 0.0165 | 0.0100 |
| **X09(3)** | 0.0013 | 0.0015 | 0.0031 | 0.0035 | 0.0022 | 0.0013 |
| **X09(4)** | 0.0013 | 0.0015 | 0.0032 | 0.0036 | 0.0033 | 0.0028 |
| **X09(5)** | 0.0017 | 0.0020 | 0.0045 | 0.0051 | 0.0041 | 0.0031 |
| **ICCG(1)** | 0.0040 | 0.0079 | 0.0446 | 0.0347 | 0.0188 | 0.0096 |
| **ICCG(2)** | 0.0031 | 0.0055 | 0.0338 | 0.0276 | 0.0139 | 0.0068 |
| **ICCG(3)** | 0.0035 | 0.0055 | 0.0202 | 0.0145 | 0.0069 | 0.0029 |
| **ICCG(4)** | 0.0026 | 0.0045 | 0.0241 | 0.0179 | 0.0088 | 0.0039 |
| **ICCG(5)** | 0.0032 | 0.0051 | 0.0263 | 0.0201 | 0.0100 | 0.0045 |
| **C08(1)** | 0.0008 | 0.0011 | 0.0018 | 0.0025 | 0.0052 | 0.0023 |
| **C08(2)** | 0.0008 | 0.0011 | 0.0019 | 0.0022 | 0.0114 | 0.0055 |
| **C08(3)** | 0.0006 | 0.0008 | 0.0013 | 0.0012 | 0.0033 | 0.0026 |
| **C08(4)** | 0.0007 | 0.0009 | 0.0013 | 0.0010 | 0.0025 | 0.0020 |
| **C08(5)** | 0.0007 | 0.0008 | 0.0011 | 0.0009 | 0.0040 | 0.0033 |
| **Control(1)** | < LOD | <LOD | 0.00014 | 0.00018 | 0.00017 | 0.00016 |
| **Control(2)** | < LOD | <LOD | 0.00015 | 0.00015 | 0.00014 | 0.00015 |
| **Control(3)** | < LOD | <LOD | 0.00013 | 0.00016 | 0.00016 | 0.00014 |

< LOQ - below the lowest analyte concentration that can be quantitatively detected with a stated accuracy and precision;
< LOD - no peak found

**Table S5.** HPLC quantification of **BHET** (mg/mL) in reaction supernatants. Reactions were carried out using 40 nM enzyme at 68°C.

| Enzyme (replicate #) | Time, h | | | | | |
|---|---|---|---|---|---|---|
| | **2** | **4** | **24** | **48** | **96** | **144** |
| **C09(1)** | 0.0001 | 0.0002 | <LOD | <LOD | <LOQ | <LOD |
| **C09(2)** | 0.0001 | 0.0003 | <LOD | <LOQ | <LOD | <LOD |
| **C09(3)** | 0.0001 | 0.0003 | <LOD | <LOD | <LOD | <LOD |
| **C09(4)** | 0.0001 | 0.0003 | <LOD | <LOD | <LOD | <LOD |
| **C09(5)** | 0.0001 | 0.0003 | <LOD | <LOD | <LOD | <LOD |
| **X05(1)** | 0.00011 | 0.0002 | <LOQ | <LOQ | <LOQ | <LOQ |
| **X05(2)** | 0.00020 | 0.0003 | <LOQ | <LOQ | <LOQ | <LOD |
| **X05(3)** | 0.00014 | 0.0002 | <LOQ | <LOQ | <LOQ | <LOQ |
| **X05(4)** | 0.00022 | 0.0004 | <LOQ | <LOQ | <LOQ | <LOQ |
| **X05(5)** | 0.00012 | 0.0002 | <LOQ | <LOQ | <LOQ | <LOD |
| **X09(1)** | 0.00092 | 0.0021 | <LOQ | <LOQ | <LOQ | <LOD |
| **X09(2)** | 0.00091 | 0.0018 | <LOQ | <LOQ | <LOQ | <LOQ |
| **X09(3)** | 0.00151 | 0.0022 | <LOQ | <LOQ | <LOQ | <LOQ |
| **X09(4)** | 0.00083 | 0.0015 | <LOQ | <LOQ | <LOQ | <LOD |
| **X09(5)** | 0.00107 | 0.0021 | <LOQ | <LOQ | <LOQ | <LOD |
| **ICCG(1)** | 0.00077 | 0.0020 | <LOQ | 0.0001 | <LOQ | <LOD |
| **ICCG(2)** | 0.00100 | 0.0010 | <LOQ | <LOQ | <LOQ | <LOD |
| **ICCG(3)** | 0.00096 | 0.0018 | <LOQ | 0.0001 | <LOQ | <LOQ |
| **ICCG(4)** | 0.00049 | 0.0016 | <LOQ | <LOQ | <LOQ | <LOD |
| **ICCG(5)** | 0.00078 | 0.0016 | <LOQ | <LOQ | <LOQ | <LOD |
| **C08(1)** | 0.00012 | 0.0003 | <LOQ | <LOQ | <LOQ | <LOQ |
| **C08(2)** | 0.00033 | 0.0006 | <LOQ | <LOQ | <LOQ | <LOQ |
| **C08(3)** | 0.00035 | 0.0007 | <LOQ | 0.0001 | <LOQ | <LOQ |
| **C08(4)** | 0.00024 | 0.0007 | <LOQ | <LOQ | <LOQ | <LOQ |
| **C08(5)** | 0.00025 | 0.0006 | <LOQ | <LOQ | <LOQ | <LOQ |
| **Control(1)** | < LOD | <LOD | < LOD | <LOD | < LOD | <LOD |
| **Control(2)** | < LOD | <LOD | < LOD | <LOD | < LOD | <LOD |
| **Control(3)** | < LOD | <LOD | < LOD | <LOD | < LOD | <LOD |

< LOQ - below the lowest analyte concentration that can be quantitatively detected with a stated accuracy and precision;
< LOD - no peak found

**Table S6.** HPLC quantification of **BHET** (mg/mL) in reaction supernatants. Reactions were carried out using 40 nM enzyme at 80 °C.

| Enzyme (replicate #) | Time, h | | | | | |
|---|---|---|---|---|---|---|
| | **2** | **4** | **24** | **48** | **96** | **144** |
| **C09(1)** | 0.0037 | 0.0050 | 0.0010 | 0.0002 | < LOQ | < LOD |
| **C09(2)** | 0.0031 | 0.0045 | 0.0008 | 0.0002 | < LOQ | < LOD |
| **C09(3)** | 0.0031 | 0.0045 | 0.0009 | 0.0001 | < LOQ | < LOD |
| **C09(4)** | 0.0028 | 0.0047 | 0.0011 | 0.0001 | < LOQ | < LOD |
| **C09(5)** | 0.0033 | 0.0039 | 0.0008 | 0.0001 | < LOQ | < LOD |
| **X05(1)** | 0.00004 | < LOD | 0.00012 | < LOD | < LOQ | < LOQ |
| **X05(2)** | < LOD | < LOD | < LOD | < LOD | < LOQ | < LOQ |
| **X05(3)** | < LOD | < LOD | < LOD | < LOD | < LOQ | < LOQ |
| **X05(4)** | < LOD | < LOD | < LOD | < LOD | < LOQ | < LOQ |
| **X05(5)** | < LOD | < LOD | < LOD | < LOD | < LOQ | < LOQ |
| **X09(1)** | < LOD | < LOD | 0.00066 | 0.00135 | 0.00035 | 0.00006 |
| **X09(2)** | < LOD | < LOD | 0.00002 | 0.00019 | 0.00026 | < LOD |
| **X09(3)** | < LOD | < LOD | < LOD | < LOD | < LOD | < LOQ |
| **X09(4)** | < LOD | < LOD | < LOD | < LOD | < LOD | < LOD |
| **X09(5)** | < LOD | < LOD | < LOD | < LOD | < LOD | < LOD |
| **ICCG(1)** | 0.00010 | 0.00032 | 0.00078 | 0.00000 | < LOD | < LOD |
| **ICCG(2)** | 0.00006 | 0.00013 | 0.00080 | 0.00002 | < LOD | < LOD |
| **ICCG(3)** | 0.00006 | 0.00014 | 0.00020 | < LOQ | < LOD | < LOD |
| **ICCG(4)** | 0.00001 | 0.00012 | 0.00033 | 0.00000 | < LOD | < LOD |
| **ICCG(5)** | 0.00005 | 0.00013 | 0.00038 | < LOQ | < LOD | < LOD |
| **C08(1)** | < LOQ | < LOQ | < LOD | < LOQ | < LOQ | < LOD |
| **C08(2)** | < LOQ | < LOQ | < LOD | < LOQ | 0.00000 | < LOQ |
| **C08(3)** | < LOQ | < LOQ | < LOD | < LOD | 0.00008 | < LOQ |
| **C08(4)** | < LOQ | < LOQ | < LOD | < LOD | 0.00001 | < LOQ |
| **C08(5)** | < LOQ | < LOQ | < LOD | < LOD | 0.00002 | < LOQ |
| **Control(1)** | < LOD | <LOD | < LOD | <LOD | < LOD | <LOD |
| **Control(2)** | < LOD | <LOD | < LOD | <LOD | < LOD | <LOD |
| **Control(3)** | < LOD | <LOD | < LOD | <LOD | < LOD | <LOD |

< LOQ - below the lowest analyte concentration that can be quantitatively detected with a stated accuracy and precision

< LOD - no peak found

**Table S7.** Diffraction data collection and refinement statistics. Statistics for the highest-resolution shell are shown in parentheses.

| Structure | C09 (DRK3) |
|---|---|
| PDB code | 8CMV |
| Wavelength (Å) | 0.7338 |
| Space group | $P6_3$ |
| Unit cell axes (Å) | a=108.876, b=108.876, c=35.402, |
| Unit cell angles (°) | α =90.00, β =90.00, γ =120.00 |
| Resolution range (Å) | 54.497-1.280 (1.302-1.280) |
| Total reflections | 1293458(62613) |
| Unique reflections | 62178(3088) |
| Multiplicity | 20.8(20.3) |
| Completeness (%) | 100.0 (100.0) |
| mean(I) / sig(I) | 6.5 (0.3) |
| Wilson B-factor | 15.27 |
| Rmerge | 0.294 (13.691) |
| Rmeas | 0.302 (14.041) |
| Rpim | 0.066 (3.103) |
| CChalf | 0.998 (0.346) |
| Reflections in refinement | 62017 (6053) |
| Reflections in free set | 3153 (317) |
| Rwork | 0.1634 |
| Rfree | 0.1871 |
| RMSD bonds (Å) | 0.012 |
| RMSD angles (°) | 1.77 |
| Ramachandran favoured (%) | 98.44 |
| Ramachandran allowed (%) | 1.56 |
| Ramachandran outliers (%) | 0 |
| Rotamer outliers (%) | 0.9 |
| Clash score | 5.65 |
| Overall number of atoms (non-H) | 2286 |
| in macromolecules | 2010 |
| in ligands | 28 |
| in solvent | 248 |
| Average B-factor(Å$^2$) | 16.14 |
| for macromolecules | 14.3 |
| for ligands | 37.33 |
| for solvent | 28.7 |
